# Cationic heme-mimetic gallium porphyrin kills bacteria and disrupts biofilms

**DOI:** 10.64898/2025.12.11.693697

**Authors:** Agata Woźniak-Pawlikowska, Klaudia Szymczak, Natalia Burzyńska, Michał Karol Pierański, Dominika Goik, Marta Kubicka, Michał Rychłowski, Lei Zhang, Joanna Nakonieczna, Mariusz Grinholc

## Abstract

The emergence of antimicrobial resistance in ESKAPE pathogens remains a clinical challenge and limits the utility of conventional antibiotics. Antimicrobial photodynamic inactivation (aPDI) using metal porphyrins is of interest, but the activity of heme-mimetic gallium porphyrins against structured biofilms has not been well defined. In this study, we evaluated a newly synthesized cationic heme-mimetic gallium porphyrin (GaCHP-2-3) activated with visible light against biofilms formed by ESKAPE representatives under static and flow conditions and on titanium surface. Light-activated GaCHP-2-3 reduced biofilm viable counts by up to 4 log_10_ CFU/mL. Prolonged serial passaging of planktonic bacteria in the presence of GaCHP-2-3 for 20 passages did not yield increases in MIC, indicating no detectable resistance development under these conditions. This work presents GaCHP-2-3 aPDI as a candidate approach for biofilm control and treatment of infections where antibiotic resistance limits the option.

## Introduction

Antibiotic resistant bacterial infections remain a major cause of significant morbidity and mortality, particularly in patients undergoing invasive procedures, immunosuppression or cancer treatment (1). Increasing global antibiotic use has accelerated the selection of multidrug-resistant strains, while the introduction of new antibiotic classes slowed substantially (2, 3).

The ESKAPE pathogens, (*Enterococcus faecium*, *Staphylococcus aureus*, *Klebsiella pneumoniae*, *Acinetobacter baumannii*, *Pseudomonas aeruginosa*, and *Enterobacter*) are major causes of healthcare-associated infections and are frequently resistant to multiple drug classes (4). Another important challenge is the effective inactivation of microorganisms growing in biofilm form. Biofilms exhibit reduced susceptibility to both host immunity and antibiotics, and are associated with chronic inflammation, tissue damage, and persistent infections (5). New non-antibiotic antibacterial modalities are therefore urgently needed (6).

Antimicrobial photodynamic inactivation (aPDI) is one such modality. In aPDI a photosensitizer is activated with visible light to generate singlet oxygen and other reactive oxygen species, leading to oxidative damage of cellular targets. Gallium compounds are of interest because Ga(III) can mimic Fe(III) and enter bacterial cells via iron uptake pathways, but cannot be reduced under physiological conditions, thereby disrupting iron-dependent function (7–9). Unlike conventional antibiotics, aPDI lethally damages microbial cells via photochemically generated ROS, and does not rely on a single molecular target.

We previously demonstrated that the heme-mimetic gallium porphyrin GaCHP accumulates intracellularly and exhibits bacterial activity against Gram-positive and Gram-negative bacteria, including multidrug-resistant strains, and that light activation enhances activity *in vitro* and *in vivo* (10, 11). GaCHP-mediated aPDI also suppressed the expression and biological activity of multiple virulence factors in *S. aureus* and *P. aeruginosa* (12)(13).

In this study, we report the synthesis and characterization of new cationic heme-mimetic gallium porphyrin, GaCHP-2-3. We show that GaCHP-2-3 aPDI kills ESKAPE pathogens, disrupts *S. aureus* and *P. aeruginosa* biofilms, and exhibits a therapeutic window in which bactericidal activity is achieved without marked host cell toxicity. We also show that serial exposure to GaCHP-2-3 aPDI produces unstable tolerance without genetic alterations. Together, these data indicate that GaCHP-2-3 represents a non-antibiotic antimicrobial modality with activity against both planktonic and biofilm-associated ESKAPE pathogens.

## Materials and Methods

### GaCHP-2-3 synthesis

Ga-CHP-2-3 was synthesized by a chelation reaction of CHP-2-3 and gallium trichloride. CHP-2-3 (100 mg, 0.127 mmol) was dissolved in anhydrous N, N-dimethylformamide (200 mL). Air in the reactor was replaced with nitrogen. An ethanol solution of gallium trichloride (25% w/w, 106 mg) was injected. The mixture was refluxed and detected with a UV–vis spectrophotometer until four bands in the Q band region of CHP-2-3 (506 nm, 538 nm, 576 nm, and 628 nm) collapsed into two bands (544 nm and 582 nm) upon metalation. The crude reaction product was dialyzed and lyophilized to obtain reddish brown powder. UV–vis (H_2_O): *λ*_max_, nm (log *ε*), 544 (4.0461), 582 (4.0526).

CHP-2-3 was synthesized according to the method we have reported (11). UV–vis (H_2_O): *λ*_max_, nm (log *ε*) 506 (4.09), 538 (4.02), 576 (3.91), 628 (3.81). ^1^H NMR (DMSO-*d*_6_, 400 MHz): *δ*, ppm 10.39, 10.32 and 10.30 (4H, 4 s, H-5, H-10, H-15 and H-20), 8.53-8.60 (2H, m, H-3^1^and H-8^1^), 8.08 (2H, t, N-H), 6.48, 6.25 (4H, m, H-3^2^ and H-8^2^), 4.39 (4H, t, H-13^1^ and H-17^1^), 3.78, 3.76, 3.68 and 3.65 (12H, 2d, H-2^1^, H-7^1^, H-12^1^ and H-18^1^), 3.18-3.17 (4H, q, H-13^4^ and H-17^4^), 3.08 (4H, t, H-13^2^ and H-17^2^), 2.42 (8H, m, H-13^5^, H-17^5^, H-13^7^ and H-17^7^), 2.24, 2.23 (12H, d, 2[N-(CH_3_)_2_]), 0.74-0.68 (4H, m, H-13^8^ and H-17^8^), 0.26 (6H, m, H-13^9^ and H-17^9^), 3.93 (2H, s, pyrrole-H). HRMS: m/z 394.2756 (calcd. for [M] ^2+^ 394.5625).

**Fig. 1.**
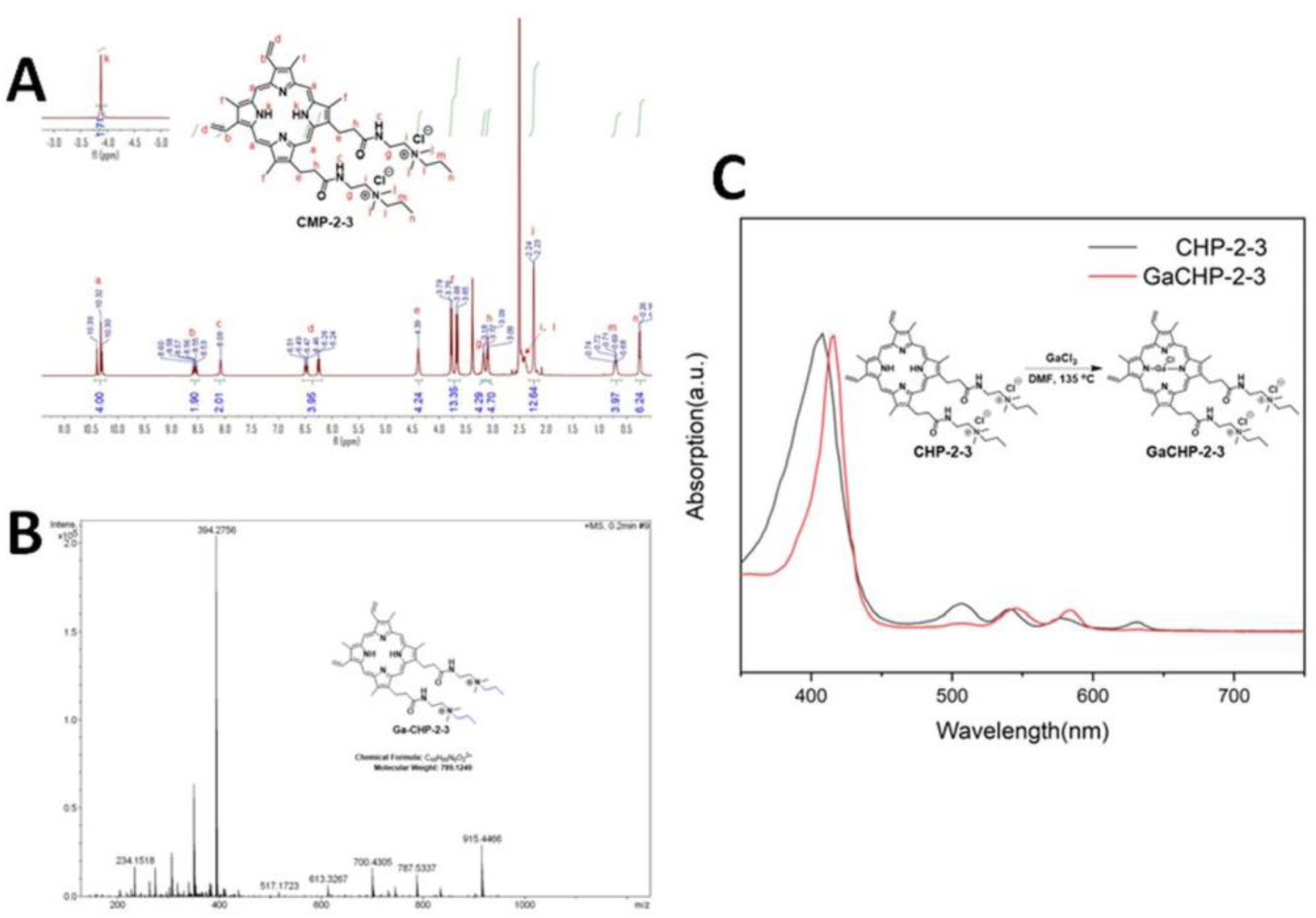
GaCHP-2-3 synthesis. Panel A) ^1^H NMR spectrum of CHP-2-3 in DMSO-*d*_6_. Panel B) Mass spectrum of CHP-2-3 obtained in the ESI^+^ mode of triple quadrupole tendem mass spectrometry. Panel C) UV-vis absorption spectra of the mixture of CHP-2-3 and gallium chloride at different complex reaction times.

### Bacteria culture

In this study, reference strains and clinical isolates of ESKAPE pathogens were grown aerobically at 37 °C with 150 rpm shaking for 16-20 h either in trypticase soy broth (TSB, bioMérieux, France) or in the case of *P. aeruginosa* in Luria-Bertani (LB) medium (Roth, Germany) (Table 1).

**Table 1.**
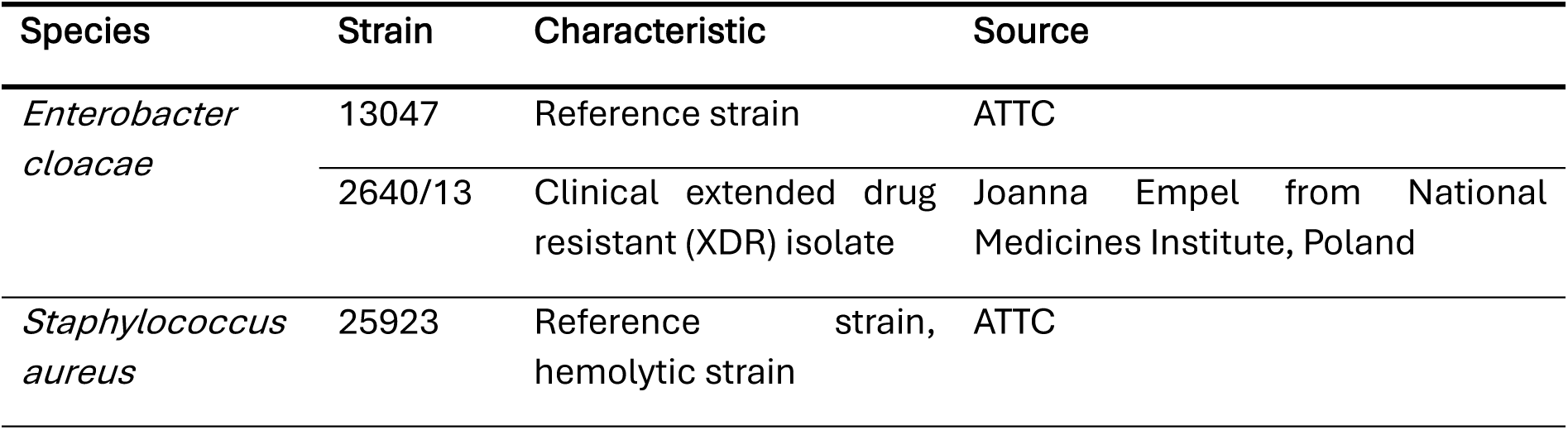
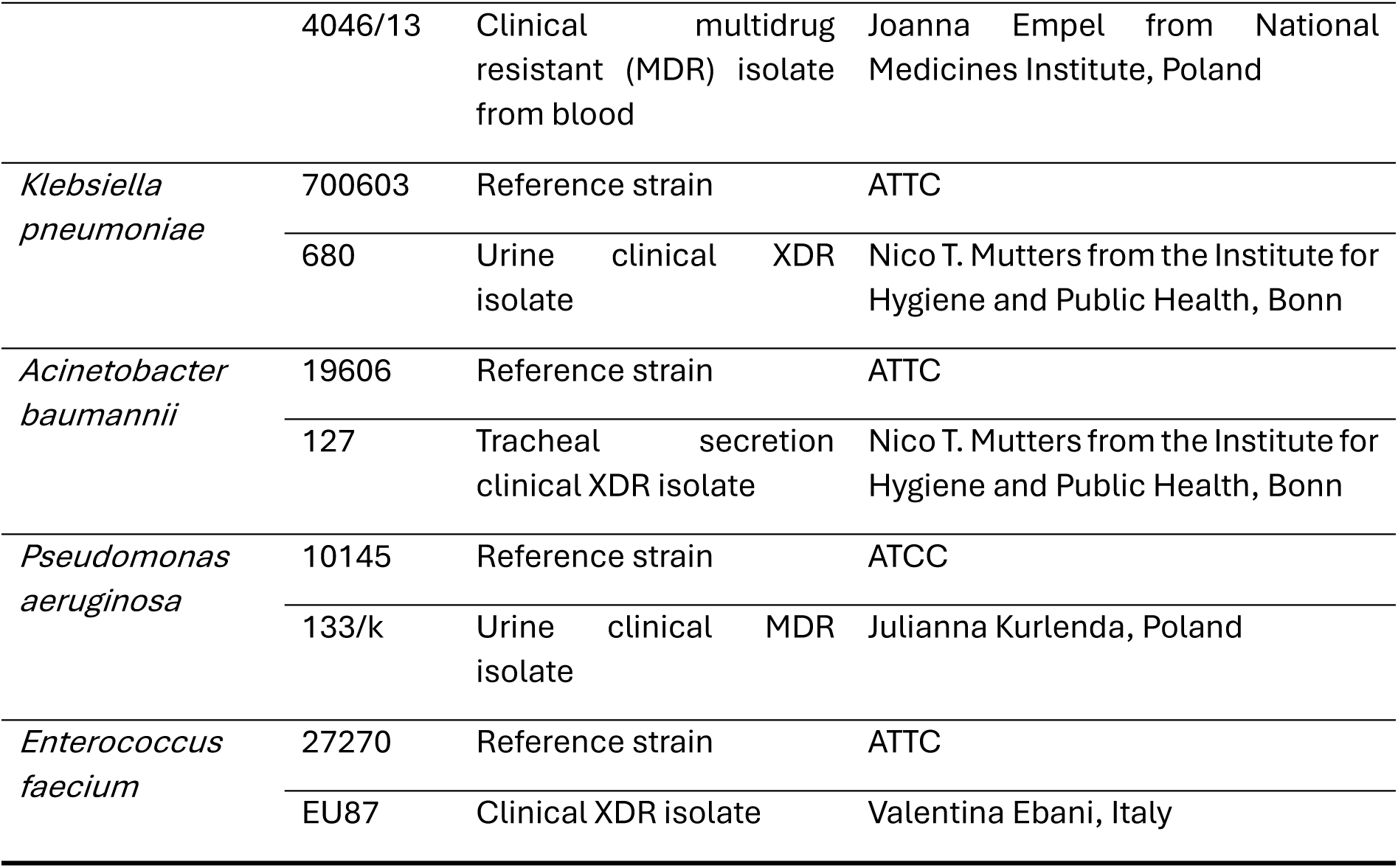
Strains used in this work.

### Cell line

HaCaT cells were cultured in DMEM supplemented with 10% of fetal bovine serum (FBS), 4.5 g/L glucose, 1 mM sodium pyruvate, 100 U/mL penicillin, 100 μg/mL streptomycin, 2 mM l-glutamine, and 1 mM nonessential amino acids. Cells were maintained at 37 °C in a 5% CO_2_ atmosphere.

### Light Source

In this study, we used a light-emitting diode (LED) light source emitting blue light (λ_max_ = 409 nm, irradiance = 5.2 mW/cm^2^) (Cezos, Poland) (14).

### Antimicrobial susceptibility testing

MBCs were determined for *S. aureus* and *P. aeruginosa.* Overnight cultures were grown in LB at 37 °C with shaking (220 rpm), diluted to 10^5^ CFU/mL in 0.3 g/L TSB. GaCHP-2-3 (1 mM stock) was 0.22-μm filtered, and two-fold diluted to 0.2 μM. In 96-well plates, 20 μL of GaCHP was mixed with 180 μL of bacterial suspension. For dark controls, plates were incubated 12 h without light; for aPDI plates were irradiated, then incubated in the dark for 12 h. CFU were enumerated on LB agar. The MBC was defined as the lowest concentration yielding ≥99.9% reduction relative to the initial control.

### Photodynamic inactivation

Overnight cultures of ESKAPE strains were diluted to 0.5 MacFarland ( ∼10^7^ CFU/mL) and transferred to a 96-well plate containing 0 - 100 μM GaCHP-2-3. Samples were incubated at 37 °C with shaking (150 rpm) for 10 min, then irradiated. Immediately after irradiation, suspensions were serially diluted in PBS and plated on TSA. CFU/mL were determined after 16-20 h incubation at 37 °C.

### Intracellular accumulation of GaCHP-2-3 in ESKAPE strains

Overnight cultures of ESKAPE strains were diluted to 0.5 McFarland and mixed with GaCHP-2-3 to final concentrations of 1, 10 or 100 µM (900 μL culture + 100 μL GaCHP-2-3) in 1.5-mL tubes. Samples were incubated at 37 °C with shaking (150 rpm) for 10 or 30 min, centrifuged (12,000 rpm; 3 min), and washed twice with PBS. Pellets were plated on TSA for CFU counting.

### Stationary biofilm studies

*S. aureus* and *P. aeruginosa* cultures were adjusted to 0.5 McFarland and 200 µL aliquots were added to a 96-well polystyrene plates. Plates were sealed to prevent evaporation, and incubated at 37 °C for 4 h (*P. aeruginosa* in LB, *S. aureus* in TSB medium). Medium was then replaced with fresh broth (200 µL), plates were resealed, and incubated at 37 °C for 20 h. For aPDI, biofilms were washed with PBS, exposed to GaCHP-2-3 (5, 10, and 25 µM) or PBS (control). Plates were incubated at 37 °C with shaking (150 rpm) for 30 min, then washed twice with PBS (100 µL).

### Flow cultivated biofilm studies

Biofilms were cultivated under flow in CDC biofilm reactor (BioSurface Technologies, Bozeman, MT) on polycarbonate coupons in BHI (*P. aeruginosa*) or TSB (*S. aureus*). After batch grown for ∼16 h at 35 °C with stirring, fresh medium was supplied by peristaltic pump for 16-20 h (10.8 mL/min) to generate mature biofilm. Coupons were transferred to PBS and exposed to GaCHP-2-3 or PBS (control) and then light. Coupons were rinsed with PBS and biofilm mass was recovered, serially diluted, plated on TSA, incubated at 37 °C for 20 h, and CFU/mL was determined.

### Titanium dental implant biofilm studies

Biofilms were established on sterilized titanium dental implants using a two-day static incubation protocol in TSB. Mature biofilm-bearing implants were washed with PBS, and exposed to GaCHP-2-3 (1, 5, and 10 µM) or PBS (control) (30 min at 37 °C, 150 rpm), and irradiated with defined light doses (up to 12.5 J/cm²). Following treatment, implants were transferred to PBS buffer, biofilm mass was disrupted by standard mechanical agitation, serially diluted, plated on TSA agar, incubated before CFU determination.

### Cyclic sublethal exposure and tolerance stability testing

*S. aureus* ATCC 25923 was subjected to repeated sublethal GaCHP-2-3 aPDI (0.2 µM, 6 J/cm^2^). Each cycle 50 µL was transferred to fresh TSB for overnight regrowth. aPDI susceptibility was reassessed after the 1^st^, 5^th^, 10^th^, and 15^th^ cycles using increasing light doses (up to 18 J/cm^2^). Tolerance stability was evaluated by serial passaging of cells from 10^th^ cycle in drug-free medium followed by re-challenge with GaCHP-2-3 aPDI. The experiment were performed in three biological replicates.

### Spontaneous mutation frequency

Spontaneous mutation frequency was determined after the 1^st^, 5^th^, 10^th^ and 15^th^ cycles of GaCHP-2-3 aPDI or ciprofloxacin (CIP) exposure. Following each cycle, cells were plated on rifampicin-containing agar and resistant colonies were quantified. Mutation frequency was determined as resistant CFU divided by total CFU of untreated controls. The experiments were performed in three independent biological replicates.

### GaCHP-2-3 accumulation

Intracellular GaCHP-2-3 (5 µM) accumulation was measured in *S. aureus* 25923 collected after the 1^st^, 5^th^, 10^th^ and 15^th^ aPDI cycles. Cells were exposed to GaCHP-2-3 under non-irradiated conditions, washed twice with PBS, lysed in 0.1 M NaOH/1% SDS (w/v), and GaCHP-2-3 fluorescence was measured in the lysate using an EnVision Multilabel Plate Reader (excitation/emission: 406 nm/582 nm). Accumulation was calculated using a fluorescence calibration curve and expressed as GaCHP-2-3 molecules per cell based on the formula shown (Fig. 2). The experiment was performed in three independent biological replicates.

### *S. aureus* growth curves after cyclic aPDI treatment

The samples from the 1^st^, 5^th^, 10^th^ and 15^th^ cycles after treatment with sub-lethal doses of aPDI (0.2 µM GaCHP-2-3, 6 J/cm^2^) were transferred to a 96-well plate (100 µL/well). Optical density (OD_600_) was measured every 30 min for 20 h at 37°C, 150 rpm in an EnVision Multilabel Plate Reader. The control group included bacterial cells non treated with light or PS, and the experiment was performed in three independent biological replicates.

### Cytotoxicity and phototoxicity testing

HaCaT cells (1 x 10 cells/well) were seeded in 96-well plates and incubated for 24 h under standard culture conditions. The next day, GaCHP-2-3 was added to the culture medium at final concentrations of 5 µM, 10 µM, and 100 µM, and the cells were incubated for 15 min. Photosensitizer-containing medium was then removed, cells were washed with Dulbecco’s phosphate-buffered saline without calcium and magnesium (DPBS—Gibco, Thermo Fisher Scientific, USA), and fresh medium was added. Cells were irradiated with blue light at final fluences ranging from 0 J/cm² to 18.72 J/cm². Cytotoxicity was assessed 24 h later using MTT assay. MTT (Invitrogen, Thermo Fisher Scientific, USA) was dissolved in DPBS (5 mg/mL), and 20 µl of this solution was added per well, followed by 4 h incubation. The MTT-containing medium was removed, and 100 µl of dimethyl sulfoxide (Thermo Fisher Scientific, USA) was added per well and gently pipetted to dissolve formazan. Absorption was measured at 550 nm after 10 s of orbital shaking at 150 rpm using an Epoch2 plate reader (BioTek, USA). Cell viability was expressed as a percentage relative to untreated, non-irradiated control. Experiments were carried out in triplicate with three technical replicates each.

### Statistical analysis

Statistical analysis was performed in Statistica 12.0 (StatSoft, Tulsa, USA) and GraphPad Prism 10.6.0 (USA). Data were expressed as mean values. To assess differences between groups, one-way or two-way ANOVA was applied, followed by post-hoc Tukey’s HSD test. In case of analyzing data without normal distribution Kruskal-Wallis analysis with Dunn’s post-hoc test was used. P values less than 0.05 were considered statistically significant.

### Production of Singlet Oxygen upon GaCHP-2-3 treatment

Singlet oxygen was quantified using Singlet Oxygen Sensor Green Fluorescent probe (SOSG; ThermoFisher Scientific) according to the manufacturer’s instructions. GaCHP-2-3 was mixed with SOSG at defined concentrations and exposed to indicated light doses (6.2 and 12.5 J/cm²). The fluorescent signal was recorded using an EnVision multiplate reader (PerkinElmer, Germany) with an excitation wavelength of 504 nm and an emission wavelength of 525 nm. Three independent experiments were performed.

### Production of total ROS upon GaCHP-2-3 treatment

Total ROS detection was quantified using the 2’,7’-dichlorofluorescein diacetate (H₂DCFDA) fluorescent probe (Invitrogen) following manufacturer instructions. GaCHP-2-3 was mixed with H₂DCFDA solution in PBS to a defined concentrations of PS (1 and 5 µM) and exposed to light doses (6.2 and 12.5 J/cm²); dark controls were included. Fluorescence was recorded using a multiplate reader (EnVision, PerkinElmer, Germany) at 495 nm/517 nm excitation/emission settings for the probe. Three independent experiments were performed.

### Ames assay

Mutagenicity was assessed using the Ames MPF Penta 10 kit (Xenometrix) with *Escherichia coli* uvrA and *Salmonella enterica* serovar Typhimurium TA98 mutant strains, following the manufacturer’s protocol. The mutant strains were exposed to GaCHP-2-3 with or without light irradiation, and to positive controls (4-nitroquinoline N-oxide for uvrA and 2-nitrofluorene for TA98). Negative controls were untreated. After incubation in indicator medium, revertants were quantified. Three independent biological experiments were performed

## Results

### Chemical modification of GaCHP increases bactericidal activity

GaCHP-2-3 was successfully synthesized through multi-step reaction. First, protoporphyrin IX (PP) was condensed with N,N-dimethylethylenediamine using oxalyl chloride as a condensation agent. The resulting tertiary amino porphyrin intermediate was then quaternized with iodopropane to yield the cation-modified porphyrin (CHP-2-3). Finally, gallium chloride was complexed with CHP-2-3, to obtain the target product, GaCHP-2-3. The successful synthesis was confirmed using ^1^H NMR and mass spectrometry, as shown in Fig. 1a and 1b. In addition, the UV-vis absorption spectra of the CHP-2-3 and its gallium derivative, GaCHP-2-3 were determined (Fig. 1c). The GaCHP-2-3 was designed and optimized for maximum efficacy. The previously described GaCHP comprised a porphyrin core, Ga^3+^ metal ion, and two arms with terminal quaternary ammonium groups (Fig. 2), which improved its solubility in water, an essential feature for clinical applications. GaCHP-2-3 was further modified by adding additional carbon atoms, increasing its hydrophobicity (Fig. 2). We conducted a detailed characterization of CHP-2-3 before it was complexed with gallium ions, including NMR and MS analyses. The molecular structure of it has been clearly confirmed. The molecular structure of CHP-2-3 after being combined with gallium ions remains unchanged. Therefore, we only used the electron absorption spectroscopy to prove the successful combination of gallium ions (11, 15). We have also attempted to characterize GaCHP-2-3 using a nuclear magnetic resonance instrument, but no signal was obtained, possibly due to the presence of metal ions.

Compared to the original compound (GaCHP), the currently studied GaCHP-2-3 exerted increased bactericidal activity under both light and dark conditions. Its antimicrobial activity was investigated against Gram-positive and Gram-negative representatives (*S. aureus* and *P. aeruginosa*, respectively) (Table 2).

**Fig. 2.**
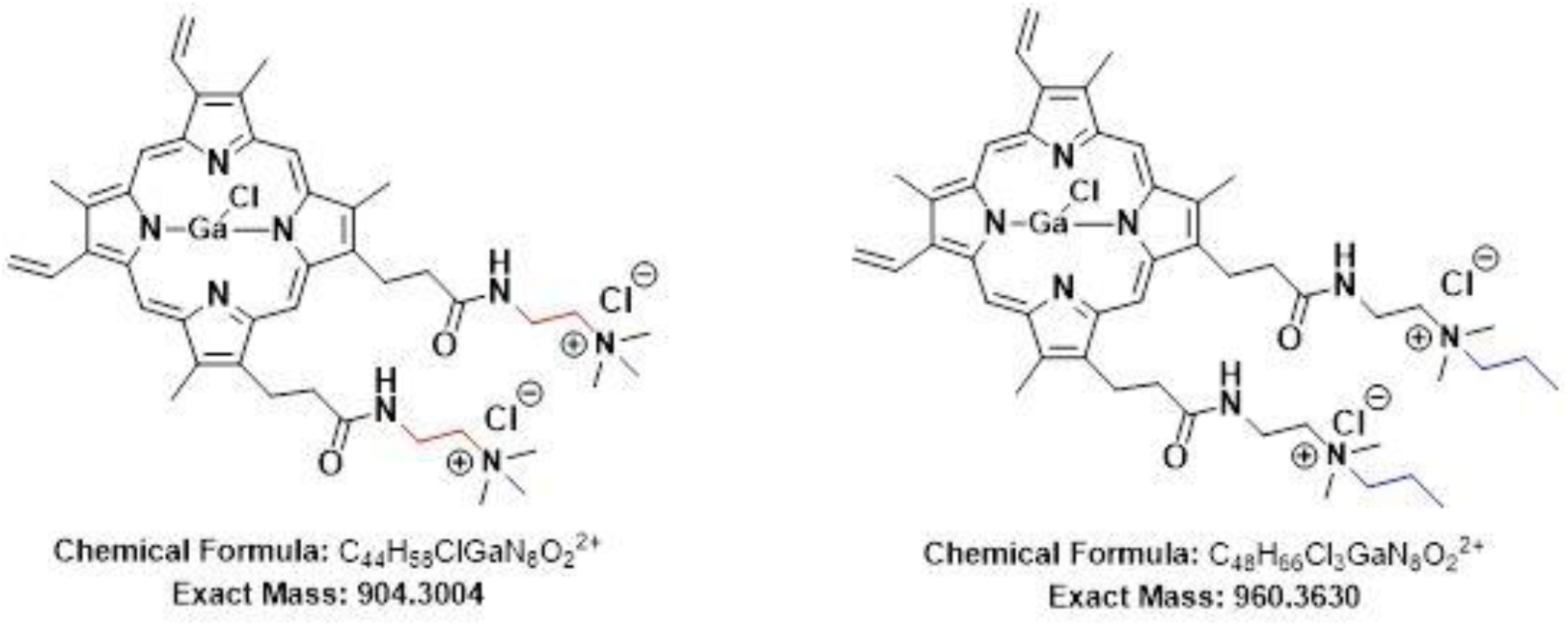
Molecular structure and molecular weight of the compounds. GaCHP (on the left) and GaCHP-2-3 (on the right).

**Table 2.** Minimum bactericidal concentrations (MBCs, μM) of gallium porphyrin compounds.

| Antibacterial agent | *S. aureus* (G+) |  | *P. aeruginosa* (G-) |  |
| --- | --- | --- | --- | --- |
|  | Dark | aPDI | Dark | aPDI |
| GaCHP | 3.1 | 1.6 | 6.3 | 6.3 |
| GaCHP-2-3 | 0.8 | 0.8 | 3.1 | 1.6 |

### GaCHP-2-3- aPDI kills ESKAPE pathogens

To evaluate broad-spectrum antibacterial activity, GaCHP-2-3 aPDI was tested against a complete set of ESKAPE pathogens, including ATCC reference strains and clinical isolates (Fig. 3). GaCHP-2-3 exhibited concentration-dependent antibacterial activity across all species. Gram-positive bacteria were more susceptible. Exposure of *S. aureus* and *E. faecium* to 1 µM GaCHP-2-3 and light (3.1 J/cm^2^) reduced viable counts by >3 log_10_ CFU/mL (Fig. 3). *E. cloacae* and *K. pneumoniae* required the highest concentration (100 µM) for detectable killing. For all other species, GaCHP-2-3 induced rapid sterilization under aPDI conditions. Together, these data show that GaCHP-2-3 aPDI is broadly active against ESKAPE pathogens, with potent activity particularly against Gram-positive bacteria.

**Fig. 3.**
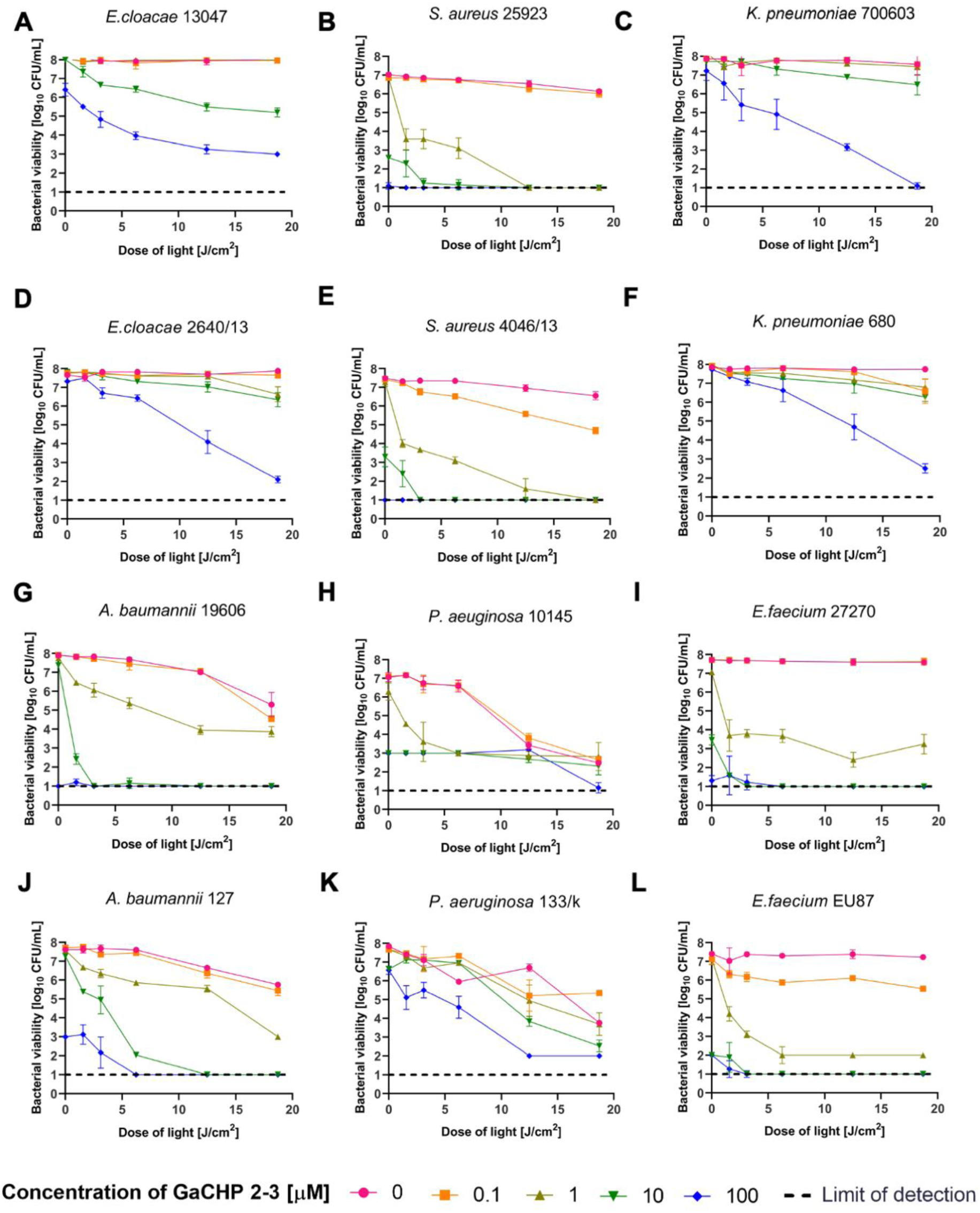
GaCHP-2-3-mediated aPDI against ESKAPE pathogens. ESKAPE representatives: reference ATCC (panels A-C, G-I) and clinical isolates (panels D-F, J-L) were exposed to various GaCHP-2-3 concentrations (0-100 µM) and divergent light doses (0-20 J/cm^2^). The bacterial viability was estimated as log_10_ of CFU/ml in accordance with non-treated samples. Values represented on the graphs are means with ±SD of each experiment performed with three biological repetitions.

### Intracellular GaCHP-2-3 accumulation correlates with aPDI efficacy

To examine determinants of susceptibility, intracellular accumulation of GaCHP-2-3 was quantified in ESKAPE reference strains (Fig. 4). GaCHP-2-3 exhibited concentration-dependent uptake in all species, and accumulation levels tracked with bactericidal activity upon light exposure. Uptake was the highest in Gram-positive bacteria and *A. baumannii*, which also corresponded to the strongest aPDI killing (Fig. 3). Despite quantitative differences among species, both Gram-positive and Gram-negative bacteria accumulated GaCHP-2-3 to levels sufficient to enable bactericidal aPDI.

**Fig. 4.**
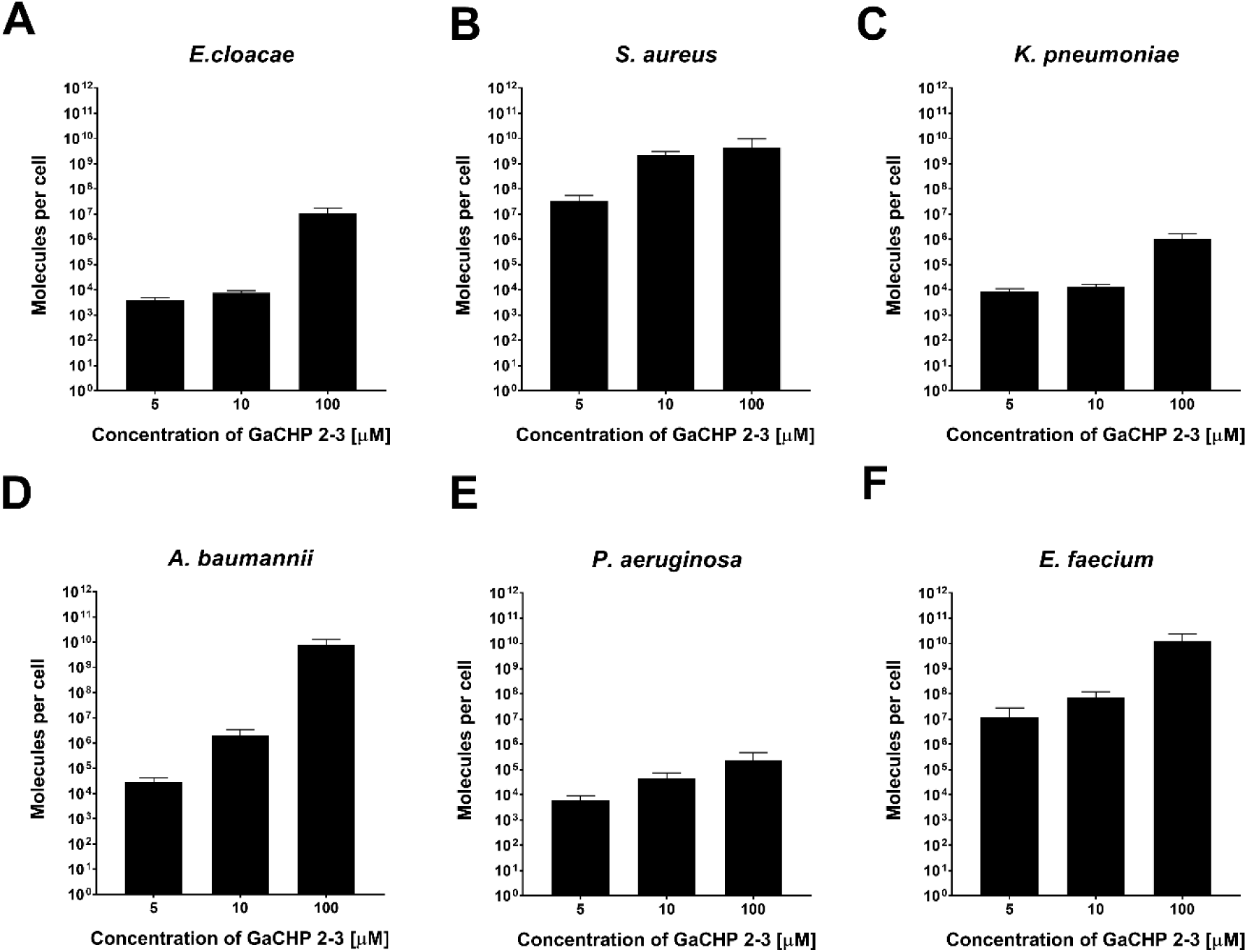
Intracellular accumulation of GaCHP-2-3. The accumulation of GaCHP-2-3 in ATCC reference strains of ESKAPE pathogens was assessed following exposure to varying concentrations of GaCHP-2-3 (5-100 µM). Accumulation is expressed as the number of molecules per cell. Data represent the means ±SD from three biological repetitions.

### GaCHP-2-3 aPDI kills of *S. aureus* and *P. aeruginosa* biofilms

GaCHP-2-3 was further evaluated in multiple biofilm models: stationary, flow-conditioned biofilms, and biofilm formed on titanium dental implants. Across models, GaCHP-2-3 exhibited concentration-dependent antibiofilm activity in both *S. aureus* and *P. aeruginosa*. In stationary biofilms 25 µM GaCHP-2-3 plus 25 J/cm^2^ reduced viability by 3.5 log_10_ CFU/mL (*S. aureus*) and 4.5 log_10_ CFU/mL (*P. aeruginosa*). Comparable reductions (3.5 and 4.0 log_10_ CFU/mL) were observed in biofilms grown under flow conditions (Fig. 5).

On titanium implant model, 10 μM GaCHP-2-3 plus 12.5 J/cm^2^ yielded the largest reduction (4.17 log_10_ CFU per implant), and 3 log_10_ CFU/implant, were achieved at 1 and 5 μM GaCHP-2-3 with 6 J/cm^2^ (Fig. 5E). Interestingly, light alone reduced viability of *S. aureus* biofilms on titanium (0.74 and 1.71 log_10_ CFU/implant at 6 and 12.5 J/cm^2^, respectively). GaCHP-2-3 in the absence of light also reduced viability on titanium (2.88 and 3.08 log_10_ CFU/ implant at 5 and 10 μM, respectively). Notably, these light-independent effects were not observed in stationary or flow-conditioned models.

**Fig. 5.**
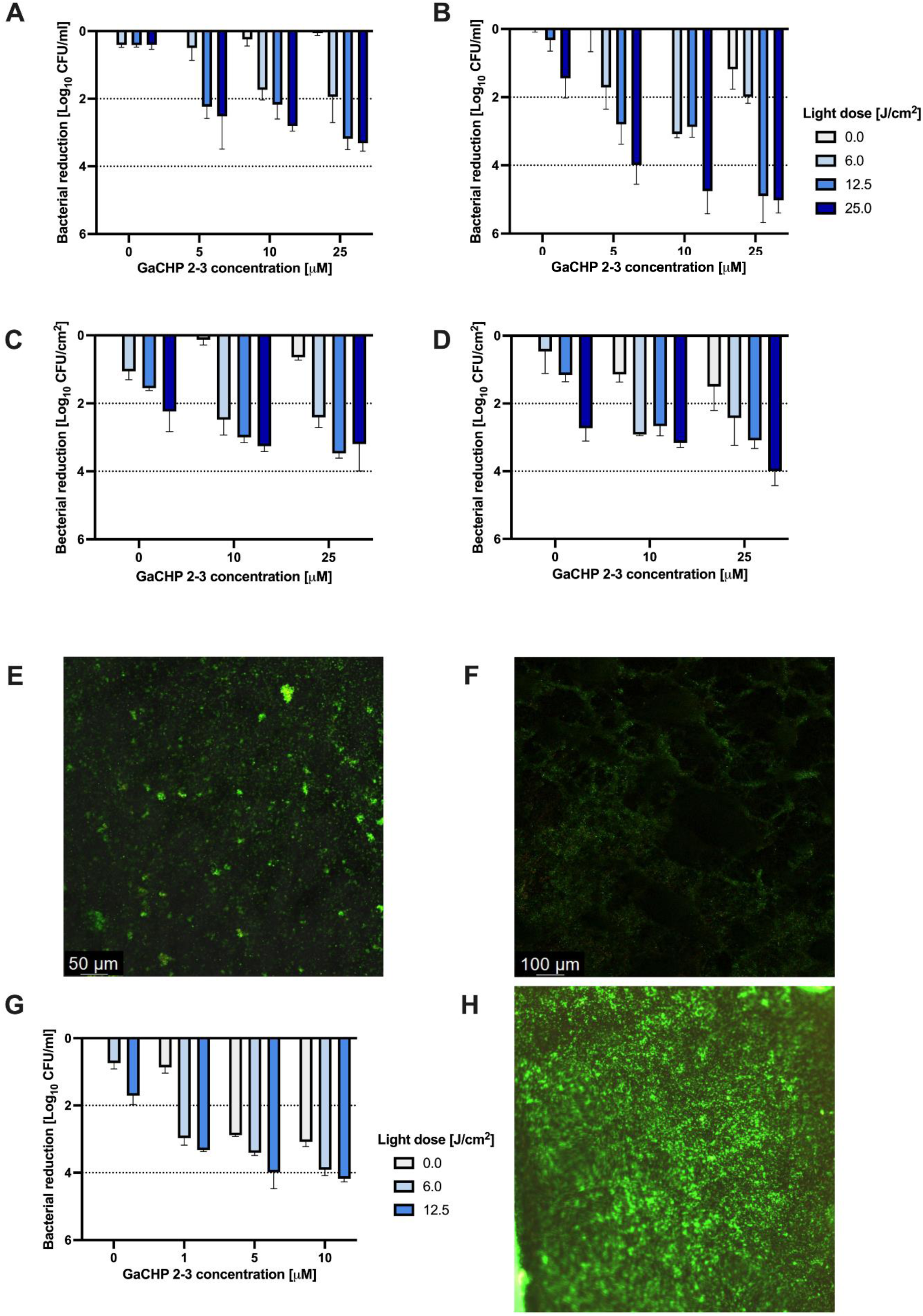
GaCHP-2-3 aPDI biofilm studies. *S. aureus* (A, C, E, G, H) and *P. aeruginosa* (B, D, F) biofilms were exposed to GaCHP-2-3 (0 - 25 µM) and light (0 - 25 J/cm^2^). (A, B) Stationary biofilm model (96-well plate). (C, D) Mature (48-h-formed) biofilms grown under flow (polycarbonate coupons, CDC bioreactor). Live-dead fluorescence image of *S. aureus* (E) and *P. aeruginosa* (F) biofilm formed on the polycarbonate coupons. (G) Bacterial viability reduction in mature (48-h-formed) biofilm on dental implants. Panel H) Live-dead fluorescence imaging of *S. aureus* biofilm on titanium implant. Data are means ±SD from three biological replicates.

### Serial GaCHP-2-3 aPDI cycles induce unstable tolerance without genetic changes

To assess the potential for resistance evolution, *S. aureus* was subjected to 15 consecutive sub-lethal cycles of GaCHP-2-3 aPDI (0.2 µM; 6 J/cm^2^) (Fig.6). Further studies concerning tolerance stability (Fig. 6E) and aPDI-induced genetic alterations supported the hypothesis that the observed tolerance was unstable and did not result from genetic alterations (Fig. 6G).

A decrease in susceptibility was detected beginning at 5^th^ cycle, with 3 log_10_ CFU/mL reduced killing relative to baseline (Fig. 6B). However, this tolerance was not stable: after five passages without aPDI, susceptibility reverted to baseline (Fig. 6E).

To probe mechanism, we tested whether tolerance was associated with increased mutation frequency. Repeated GaCHP-2-3 aPDI did not increase rifampicin-resistant mutant frequency (Fig.6G). We next quantified GaCHP-2-3 accumulation in cells sampled from the 1^st^, 5^th^, 10^th^ and 15^th^ cycles. GaCHP-2-3 uptake per *S. aureus* cell was unchanged across cycles (Fig. 6H). Thus, tolerance was transient, not genetic, and not explained by photosensitizer accumulation.

**Fig. 6.**
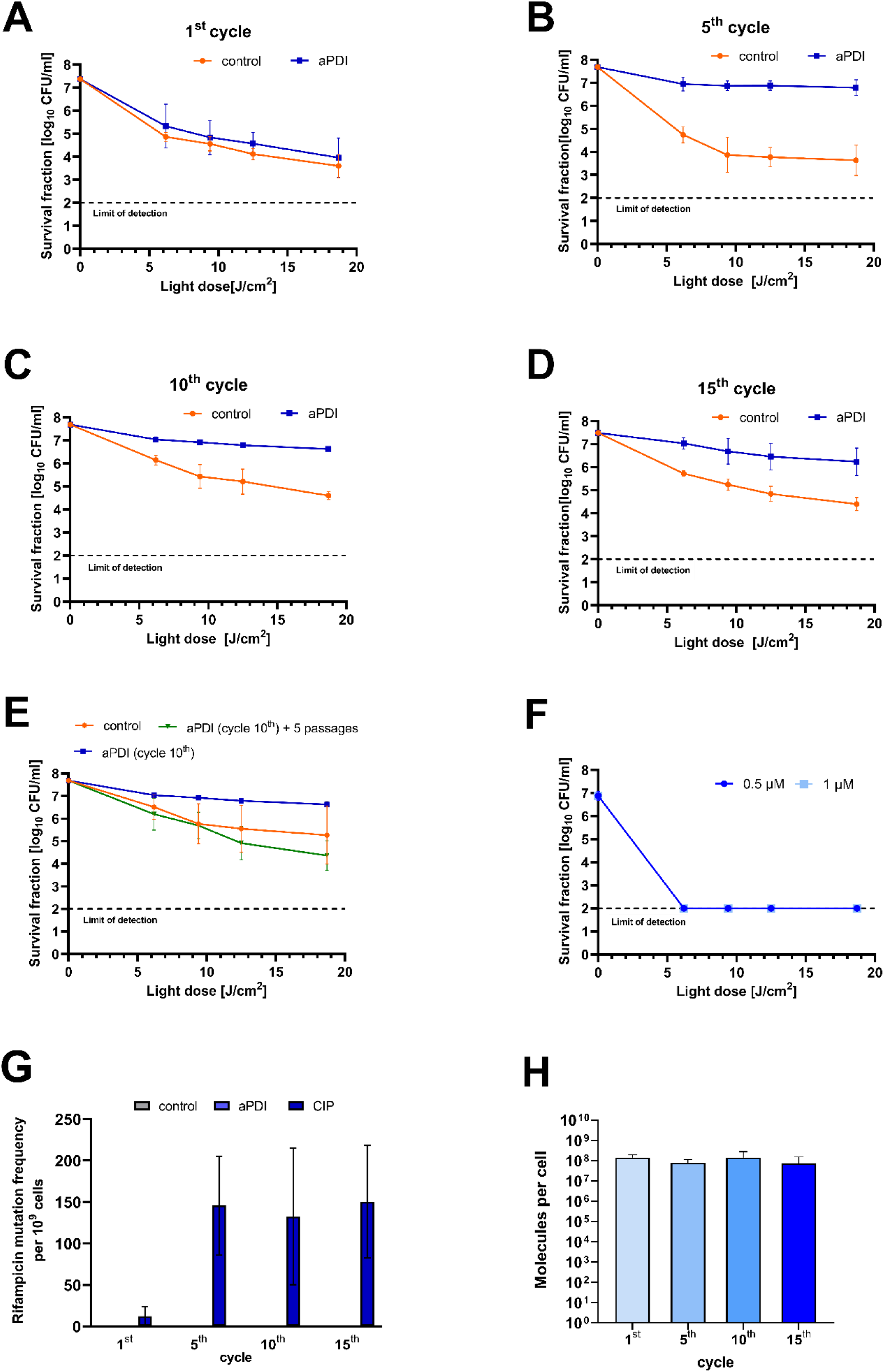
***S. aureus* tolerance development.** (A-D) The susceptibility of *S. aureus* to 0.2 µM GaCHP-2-3 aPDI after 1^st^, 5^th^, 10^th^ and 15^th^ sub-lethal cycles. (E) Tolerance stability: cells from 10^th^ cycle after 5 passages without selection. (F) The susceptibility of 10^th^ cycle *S. aureus* to 0.5 and 1 µM GaCHP-2-3 aPDI. (G) Mutation frequency after sub-lethal GaCHP-2-3 aPDI vs ciprofloxacin (CIP). (H) Intracellular GaCHP-2-3 accumulation in *S. aureus* from 1^st^,5^th^,10^th^ and 15^th^ cycle. Data are means ± SD (three biological replicates).

### Sub-lethal GaCHP-2-3 aPDI alters *S. aureus* growth dynamics

To evaluate whether the tolerance was associated with altered growth dynamics, growth curves of *S. aureus* from 1^st^, 5^th^, 10^th^ and 15^th^ cycles were compared to untreated *S. aureus* (Fig. 7). Absorbance measurements (OD_600_) revealed a progressive delay in entry into the logarithmic growth phase, beginning at the 1^st^ cycle and increasing with subsequent cycles. Quantitative growth parameters (μ_max_, *T*_d_, time to stationary phase, and *A*_max_) are shown in Table 3. Relative to untreated cells, μ_max_ decreased and *T*_d_ increased across all cycles. Further analysis of the time required to reach stationary phase supported this observation. Time to stationary phase increased from 450 min in untreated cells to 540, 660, 720 and 870 min in 1^st^, 5^th^, 10^th^ and 15^th^ cycles, respectively. At 450 min (the end time of exponential phase in untreated *S. aureus*), relative growth declined from 44.5 at the 1^st^ cycle to 17.13% at the 15^th^ cycle (Table 3). Thus, repeated sub-lethal GaCHP-2-3 aPDI cycles progressively reduced the growth capacity of *S. aureus*.

**Fig. 7.**
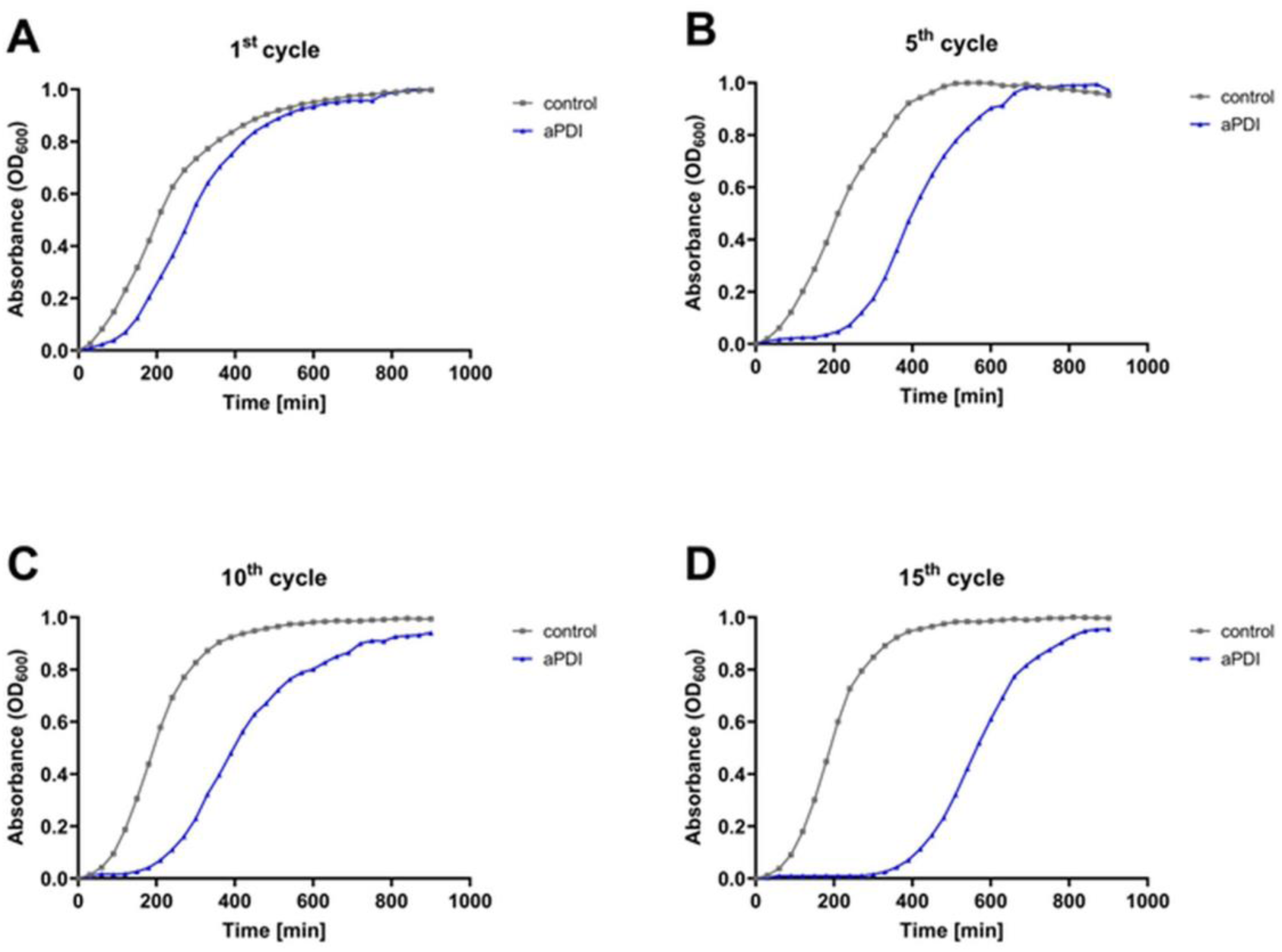
Effect of GaCHP-2-3 aPDI sub-lethal phototreatment on *S. aureus* growth dynamics. Panels A,B,C and D) Normalized growth curves of *S. aureus* after aPDI treatment in 1^st^ (A), 5^th^ (B), 10^th^ (C) and 15^th^ (D) cycle.

**Table 3.** Staphylococcal growth parameters upon consecutive exposures to aPDI.

| Treatment | Parameters of *S. aureus* growth |  |  |  |  |
| --- | --- | --- | --- | --- | --- |
| | $\mu_{\max}$<br>[OD<sub>600</sub>/h] | Td<br>[hours] | $A_{\max}$<br>[OD<sub>600max</sub>] | Time of<br>obtained<br>stationary<br>phase [min] | *S. aureus* growth<br>at the end of<br>exponential phase<br>[%] |
| Untreated | 0.279 | 2.48 | 0.688 | 450 | 100 |
| aPDI 1<sup>st</sup> cycle | 0.193 | 3.59 | 0.447 | 540 | 44.50 |
| aPDI 5<sup>th</sup> cycle | 0.210 | 3.30 | 0.507 | 660 | 21.97 |
| aPDI 10<sup>th</sup> cycle | 0.168 | 4.13 | 0.436 | 720 | 25.69 |
| aPDI 15<sup>th</sup> cycle | 0.195 | 3.55 | 0.555 | 870 | 17.13 |

$\mu_{\max}$ – maximum specific growth rate during exponential phase of bacterial growth; Td – time of duplication, also known as generation time; $A_{\max}$ – maximal absorbance value with maximal bacterial density; Time of the obtained stationary phase – defined as the point of curve flattening; *S. aureus* growth at the end of exponential phase [%] at 240 min as the inflection point of the exponential growth curve calculated in the reference to control (untreated) cells (100%)

### GaCHP-2-3 activity depends on ROS generation

To test whether GaCHP-2-3 aPDI generates ROS, total ROS and singlet oxygen were quantified using DCFDA and SOSG probes, respectively (Fig. 8). Total ROS increased in a dose-dependent manner as a function of PS concentration and light dose (Fig. 8A). Singlet oxygen was detected at high light dose even without the GaCHP-2-3, but levels were markedly higher in the presence of photosensitizer (Fig. 8B). These data demonstrate that light-excited GaCHP-2-3 generates ROS, including singlet oxygen (Fig. 8B).

**Fig. 8.**
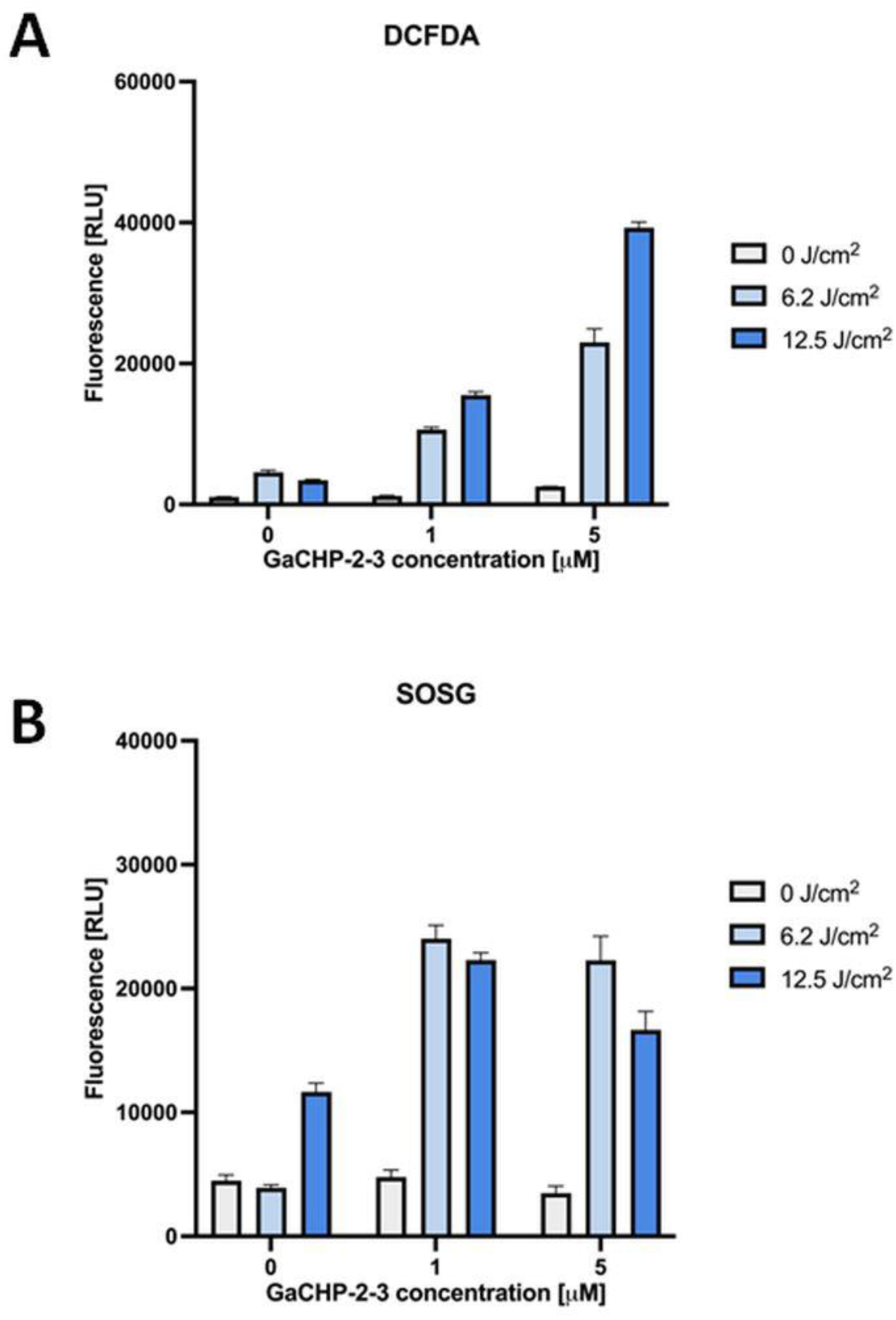
ROS generation upon GaCHP-2-3 treatment. Panel A) Assessment of the production of total ROS using the 2’,7’-dichlorofluorescein diacetate (H₂DCFDA) probe. The samples were exposed to light doses of 6.2 and 12.5 J/cm², and the level of ROS was subsequently measured at excitation and emission wavelengths of 495 and 517 nm. Panel B) Singlet oxygen production determined using a Singlet Oxygen Sensor Green probe. Samples were exposed to light doses of 6.2 and 12.5 J/cm², and then the fluorescent signals were recorded at excitation wavelength of 504 nm and an emission wavelength of 525 nm. Values represented on the graphs are means with ±SD of each experiment performed with three repetitions.

### GaCHP-2-3 shows no detectable mutagenicity

To assess mutagenic potential, GaCHP-2-3 was tested in the Ames assay under dark and light conditions using *S.* Typhimurium TA98 (Fig. 9A) and *E. coli* uvrA (Fig. 9B). GaCHP-2-3 did not increase revertant frequency in either strain. Only the positive control compounds (2-nitrofluorene (2-NF) for TA98 and 4-nitroquinoline N-oxide (4-NQO) for uvrA) produced the expected increase in revertants.

**Fig. 9.**
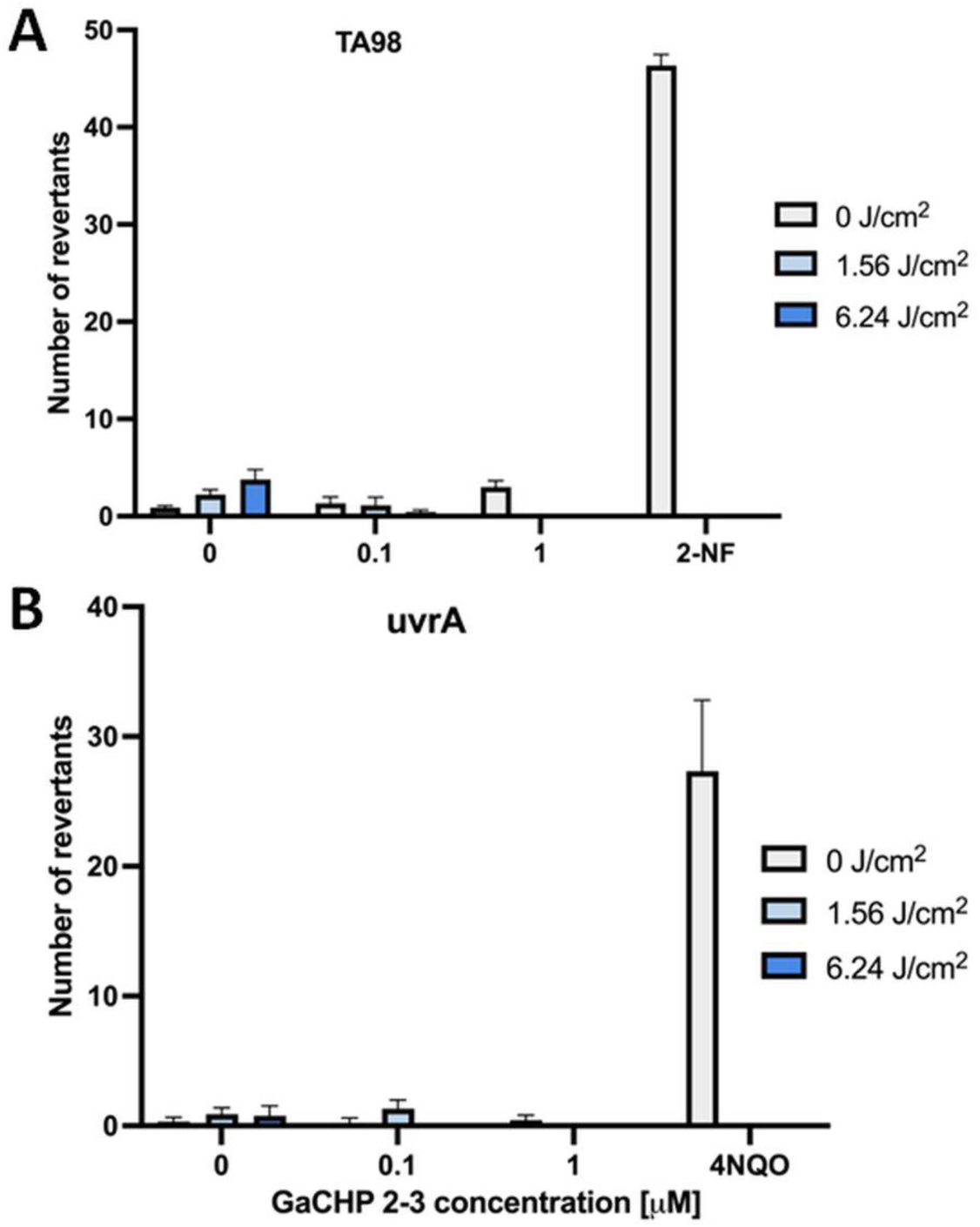
Assessment of dark and light-induced mutagenicity of GaCHP-2-3. The mutagenic effect was assessed in Ames test with two different mutants, i.e., *S.* Typhimurium TA98 (panel A) and *E. coli* uvrA (panel B). The assay included two light doses (1.56 and 6.24 J/cm^2^), two PS concentrations (0.1 and 1 µM) and positive control (4-nitroquinoline N-oxide (4-NQO) for uvrA and 2-nitrofluorene (2-NF) for TA98). The experiment was performed in three biological replicates, each replicating in three technical repetitions. Values represented on the graphs are means with ±SD of each experiment.

### GaCHP-2-3 is non-cytotoxic in the dark but induces fluence- and concentration-dependent phototoxicity

GaCHP-2-3 cytotoxicity was assessed in HaCaT keratinocytes using MTT assay, and phototoxicity was examined after 405-nm LED irradiation. In the absence of light, GaCHP-2-3 did not reduce viability at concentration up to 100 µM, indicating no detectable dark toxicity (Fig. 10). In contrast, 405-nm irradiation in the presence of GaCHP-2-3 produced concentration- and fluence-dependent reductions in viability.

To determine whether GaCHP-2-3 aPDI operates within a therapeutic window, we compared the fluences that induced killing of bacteria with those that caused phototoxicity in HaCaT cells. Fluences around 6 J/cm² were sufficient to reduce biofilm-associated pathogens by ∼2 log₁₀ CFU (99% of initial bacterial count), whereas the same fluence range produced minimal loss in HaCaT viability at concentrations ≤10 μM. Significant HaCaT phototoxicity was observed only under the highest test condition. Thus, the fluence–concentration combinations that produced bactericidal activity aligned with conditions that remained well tolerated by mammalian cells. These data establish a therapeutic window where GaCHP-2-3 aPDI is bactericidal but not cytotoxic.

**Fig. 10.**
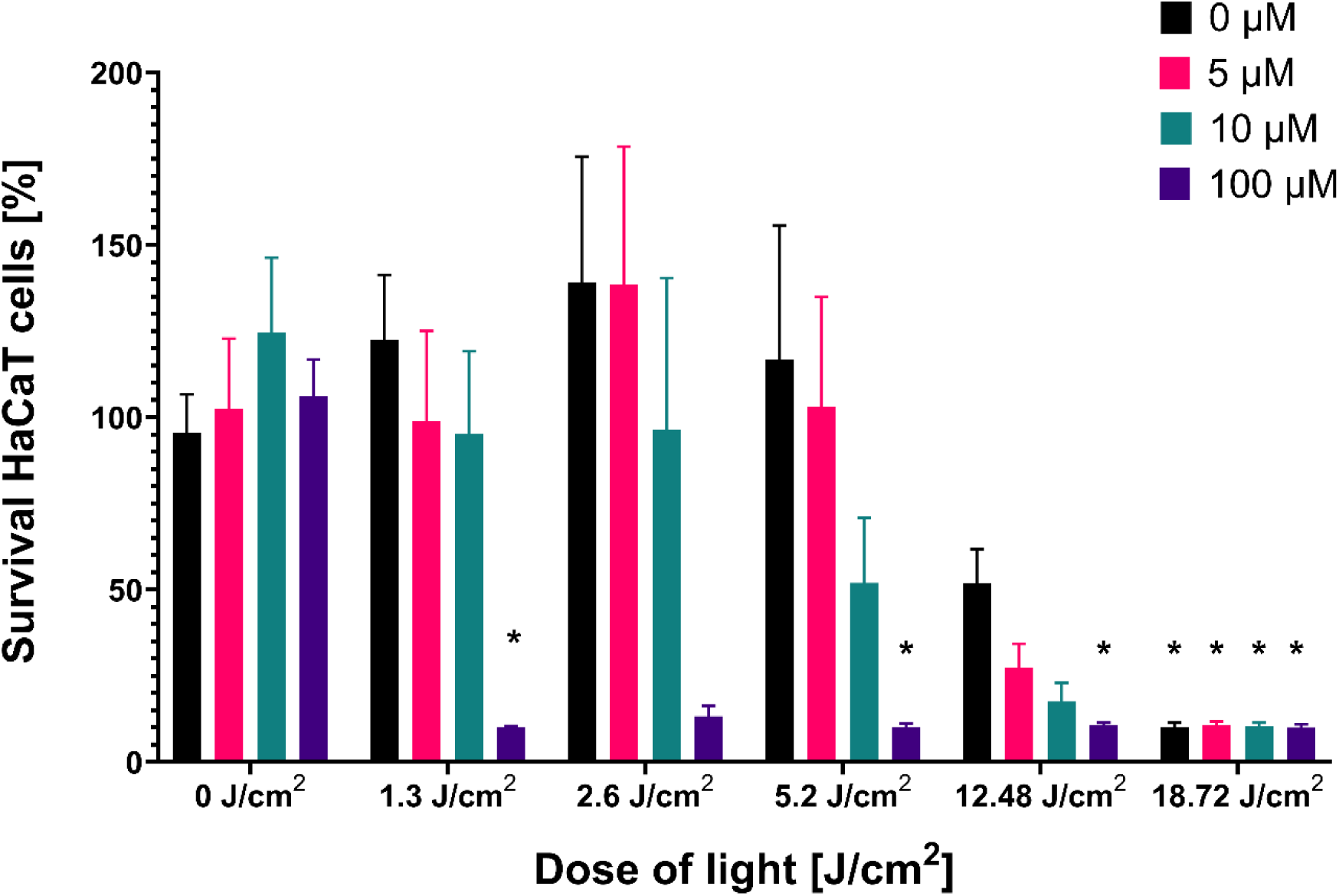
Cell viability of HaCaT cells assessed by MTT assay following treatment with the photosensitizer (0–100 μM) either in the dark or after irradiation at light doses ranging from 1.3 to **18.7 J/cm².** Data are shown as mean ± SD (n = 9). Statistical analysis was performed using the Kruskal–Wallis test followed by Dunn’s post-hoc multiple comparison test. All treatment groups were compared to the untreated, non-irradiated control (0 μM, 0 J/cm²). (* adjusted p < 0.01).*

## Discussion

Gallium porphyrins (GaPP) have been utilized as photosensitizers in antimicrobial photodynamic inactivation (aPDI), demonstrating antimicrobial effects through a complex mechanism of action (16, 17). This mechanism includes the production of reactive oxygen species (ROS) upon light activation, and disruption of iron metabolism (11, 18). When GaPP is exposed to light at a specific wavelength, it absorbs photons, transitioning from its ground state to an excited singlet state. This singlet state can undergo intersystem crossing to a more reactive, longer-lived triplet state. In this triplet state, GaPP transfers energy to molecular oxygen (O₂), generating ROS such as singlet oxygen (¹O₂), superoxide anions (O₂•⁻), hydroxyl radicals (•OH), and hydrogen peroxide (H₂O₂). These ROS are highly reactive and inflict significant damage on microbial cells. They indiscriminately oxidize cellular components, leading to various types of damage: lipid peroxidation leading to increased permeability of cell membranes and eventual cell lysis, protein oxidation resulting in its denaturation and loss of function, and nucleic acid damage including breaks in DNA and RNA strands and alter bases, impairing replication and transcription. The cumulative damage to these critical cellular components ultimately results in the death of microorganisms, making GaPP a potent antimicrobial agent (19, 20).

Gallium, also plays a critical role due to its chemical similarity to iron (Fe), though it differs in key properties. Microorganisms can mistakenly incorporate gallium into their iron-dependent processes, leading to significant disruptions in cellular function. Gallium can replace iron in iron-dependent enzymes, rendering these enzymes non-functional since gallium cannot undergo the same redox reactions as iron in physiological conditions. This substitution inhibits essential processes in microbial metabolism (11, 21). Additionally, gallium competes with iron in microbial iron acquisition systems, blocking the microbes from obtaining the iron necessary for survival and growth. This competition disrupts iron homeostasis, creating a state of iron starvation that weakens the microorganism’s defenses and increases its susceptibility to oxidative stress (22).

Although gallium porphyrins are highly potent, research on their action under light activation remains limited. Morales et al. demonstrated that Ga-protoporphyrin IX chloride (Ga-PpIX) is an effective photosensitizer against *S. aureus*, including multidrug-resistant MRSA strains (23). In their study, Ga-PpIX exhibited enhanced antimicrobial activity compared to pure protoporphyrin IX and showed lower cytotoxicity toward human keratinocytes. The same research group in further studies developed a nanoscale agent for targeted aPDI, which consisted of Ga-PpIX encapsulated in hemoglobin and assembled on silver nanoparticles (24). This innovative formulation achieved a reduction in *S. aureus* viable count by over 6 log_10_ units. The microbial burden reduction observed with this nanostructure was significantly greater, by several orders of magnitude, than with monotreatment using free Ga-PpIX.

Additionally, Awad et al. described the development of another nanostructure loaded with gallium protoporphyrin (16). This formulation was specifically designed to target microbial biofilms, which are a major challenge in aPDI due to the difficulty many photosensitizers face in penetrating the biofilm matrix. To address this limitation, liquid crystal lipid nanoparticles (LCNPs) were employed as a biocompatible delivery system for gallium protoporphyrin. This approach significantly enhanced the antibacterial activity of the photosensitizer, achieving a two-fold increase in the killing of both wild-type and MRSA biofilm cultures.

Beginning in 2022, our research series highlighted the potent antimicrobial photodynamic activity of novel gallium porphyrin derivatives. Michalska et al. (2022) demonstrated that gallium mesoporphyrin IX (Ga-MP) exhibits antimicrobial efficacy through a dual mechanism: disruption of iron metabolism and photosensitizing properties in aPDI (22). This compound achieved efficient bacterial photoinactivation, reducing *S. aureus* by over 5 log_10_ units in CFU/mL while showing relatively low cyto- and phototoxicity toward eukaryotic cells. Building on these findings, Zhang et al. proposed a novel cationic heme-mimetic gallium porphyrin derivative (GaCHP) with the addition of two quaternary ammonium groups compared to Ga-MP, further enhancing its properties (11). GaCHP was shown to rapidly eliminate multidrug-resistant and wild-type Gram-positive and Gram-negative bacterial species. Unlike antibiotics, bacteria demonstrated significant difficulty evolving resistance to GaCHP treatment. Furthermore, GaCHP was found to induce heme-assisted catalase inactivation. In a proof-of-concept study, GaCHP was tested on mice with full-thickness *S. aureus* skin infections. The infection models were treated with GaCHP administered via tail vein injection. This treatment led to significantly faster wound healing compared to the control group, with the GaCHP-treated wounds showing even greater therapeutic effectiveness than vancomycin (10, 11). The inclusion of two additional quaternary ammonium groups in GaCHP significantly enhanced its water solubility, a critical characteristic for the development of new photosensitizing agents, as it directly influences their *in vivo* antimicrobial efficacy and biocompatibility within host tissues. Our further studies by Szymczak et al. (2024) demonstrated that light-activated GaCHP not only significantly reduced microbial viability but also suppressed both the expression and biological activity of multiple virulence factors in Gram-positive and Gram-negative bacteria, including *S. aureus* and *P. aeruginosa* (12).

This study is the first to demonstrate the bactericidal effect of gallium porphyrin sensitizers using various microbial biofilm models, including stationary, flow-conditioned and titanium dental implant-associated biofilms of both, Gram-positive and Gram-negative representatives (*S. aureus* and *P. aeruginosa*, respectively). In this study, we further optimized the previously described gallium porphyrin photosensitizer, GaCHP by incorporating two additional hydrophobic carbon atoms. This modification was designed to tune the local hydrophobicity of the molecule, thereby improving its microbial binding affinity and bactericidal efficacy. As evidenced in Table 2, this chemical modification resulted in a decreased MIC for GaCHP-2-3 compared to the parental GaCHP compound, both, in the dark and upon light excitation. The newly synthesized compound displayed a significant bactericidal effect against a broad range of Gram-positive and Gram-negative pathogens, assigned to so called ESKAPE human high priority pathogens (25). Uptake studies evidenced that GaCHP-2-3 demonstrated concentration-dependent accumulation across all tested bacterial species (Fig. 4), which directly correlated with its bactericidal efficacy upon light treatment. The highest accumulation of GaCHP-2-3 was observed in Gram-positive bacteria and *A. baumannii*, corresponding to the most pronounced aPDI outcomes (Fig. 3). Despite differences in accumulation levels among species, it is noteworthy that both Gram-positive and Gram-negative bacteria effectively accumulated the compound, leading to a robust bactericidal response under light treatment. The broad-spectrum antimicrobial activity of GaCHP-2-3 was further demonstrated in various biofilm models, including stationary and flow-conditioned growth systems, as well as dental implant-associated biofilms for both Gram-positive and Gram-negative representatives. As depicted in Fig. 5, GaCHP-2-3 showed concentration-dependent antibiofilm activity across all tested bacterial species, regardless of the biofilm model. Notably, GaCHP-2-3 aPDI achieved significant antibiofilm effects against both Gram-positive and Gram-negative species, including biofilms grown under stationary conditions and mature biofilms established in dynamic flow environments (Fig. 5). Next, a key focus of this research was assessing bacterial resistance to GaCHP-2-3 following repeated treatments. While *S. aureus* subjected to up to 15 consecutive cycles of sub-lethal GaCHP-2-3 aPDI displayed detectable tolerance to the treatment, no evidence of resistance was observed (Fig. 6F).

Collectively, GaCHP-2-3 revealed excellent antibacterial activity against both planktonic and biofilm cultures of Gram-positive, Gram-negative, and multidrug-resistant strains. Importantly, it did not induce microbial resistance following the treatment. Moreover, this research confirmed that the newly synthesized compound (GaCHP-2-3) demonstrated higher bactericidal efficacy compared to its precursor GaCHP and effectively accumulated in all tested ESKAPE species. Finally, GaCHP-2-3 exhibited strong activity against mature Gram-positive and Gram-negative biofilms, indicating its great potential as an experimental therapeutic.

## Acknowledgments

The authors acknowledge the financial support from the National Science Centre in Poland (grant no. 2018/30/Q/NZ7/00281) (M.G.), Natural Science Foundation of China (grant no. 21961132005) (L.Z.). We would like to thank Magdalena Król for carrying out experiments related to the elimination of biofilm formed on dental implants.

## Declaration of interests

Authors declare no conflict of interests.

